# Neonatal phlebotomy-induced anemia compromises mitochondrial bioenergetics in the developing hippocampus

**DOI:** 10.64898/2025.12.05.692675

**Authors:** Thomas W. Bastian, Diana J. Wallin, Amanda K. Barks, Raghavendra B Rao, Michael K. Georgieff

## Abstract

**Background:** Anemia is a common medical condition in preterm infants. Previous studies show that neurodevelopmental outcomes of preterm infants are dependent in part on the degree of anemia. In a developmentally appropriately-timed neonatal mouse model, phlebotomy induced anemia (PIA) of the degree commonly seen in hospitalized preterm infants results in brain iron deficiency and hypoxia and significant short-term and long-term brain dysfunction, especially in the hippocampus. Iron and oxygen are critical for mitochondrial oxidative phosphorylation-mediated ATP production and thus energetically demanding brain developmental processes (e.g., axon/dendrite growth, myelination, synaptogenesis).

**Objective:** To test the hypothesis that neonatal PIA acutely impairs mitochondrial respiratory capacity and electron transport chain (ETC) complex function in the developing hippocampus.

**Methods:** Neonatal mice were phlebotomized daily beginning on postnatal day 3 (P3). On P14, mitochondria were isolated from the hippocampus of male and female PIA and non-bled mice. Seahorse bioenergetic analyses were performed to determine the effects of PIA on mitochondrial oxidative phosphorylation activity and ETC complex functional capacity.

**Results:** PIA hippocampal mitochondria demonstrated an overall reduced oxygen consumption rate (OCR) compared to non-bled controls when ETC oxygen consumption was coupled to ATP production. PIA reduced hippocampal mitochondrial OCR that was not due to the ETC in females not males. Basal respiration, proton leak, and maximal respiratory capacity were significantly reduced in PIA hippocampal mitochondria, an effect that did not differ by sex. When mitochondrial ETC oxygen consumption was uncoupled from ATP production with the protonophore FCCP, a mild reduction in OCR was observed across all ETC complexes, with only complex I-mediated OCR being significantly lower than non-bled controls.

**Conclusions:** These findings suggest that impaired mitochondrial energetic capacity may mechanistically contribute to the persistent neurobehavioral deficits caused by PIA, through dysregulation of energy-demanding neurodevelopmental processes (e.g., neuron structural maturation).

## Introduction

Preterm birth increases the risk of impaired neurodevelopment and lifelong neurobehavioral disabilities compared to term birth (1). While many factors influence neurodevelopment in infants born preterm, one highly controllable factor is the management of anemia of prematurity (2–9). Anemia is a common co-morbidity in premature infants with phlebotomy-induced blood loss being the major contributor (5–9). There is controversy as to whether anemia or its treatment with red cell transfusions alters neurodevelopmental outcomes in preterm infants (3,4,9–14). However, animal models of phlebotomy-induced anemia (PIA) demonstrate striking effects on brain gene expression and behavioral outcomes (9,15–18).

Thus, a better understanding of the mechanisms underlying the brain developmental impairments caused by neonatal anemia is required to inform and improve treatment management.

Brain development is a metabolically and energetically demanding process with the newborn human brain accounting for 60% of total body oxygen consumption (19,20). In addition to oxygen and macronutrients, it is critical to maintain sufficient levels of other nutrients that support cellular energy metabolism, such as iron. Mitochondrial oxidative phosphorylation (OXPHOS) is the most efficient cellular mechanism for producing cellular energy in the form of ATP. Cytochrome c and all four complexes of the electron transport chain (ETC) require iron for their ATP generating activity through iron-sulfur proteins and/or heme-containing cytochromes (21). Fetal-neonatal dietary iron deficiency anemia (IDA) decreases cytochrome c oxidase (complex IV) activity – which requires both oxygen and iron – and simplifies dendritic structure in the neonatal rat hippocampus (22–25). Iron deficiency specifically within the developing hippocampal neuron impairs mitochondrial OXPHOS, ATP production and structural development, which persist into the mature neuron without timely neuronal iron repletion (26–29).

Our mouse model of neonatal phlebotomy-induced anemia (PIA) reduces the hematocrit to levels proportionate to those tolerated in preterm infants during a time period that is developmentally equivalent to 26 to 40 weeks post-conceptional age in humans (9,15–18).

Similar to dietary IDA mouse and rat models, our mouse model of neonatal phlebotomy-induced anemia (PIA) causes a 40% decrease in overall brain iron concentration due to loss of total body iron (15). This places the neonatal PIA brain, and in particular the rapidly developing hippocampus, at risk of mitochondrial energetic dysfunction from both iron deficiency (ID) and tissue hypoxia (15,21). Neonatal mouse PIA increases lactate concentrations specifically within the postnatal day (P) 14 hippocampus (15). We speculate this is due to an increased reliance on anaerobic glycolysis, a less efficient mechanism for energy production, suggesting mitochondrial compromise due to hippocampal ID (16,30). In support of this, PIA causes increased hippocampal activity of AMPK, a master regulator of cellular energy homeostasis that is activated when cellular energy demand outstrips energy production causing a decrease in the ratio of ATP to AMP (16). Ultimately, PIA causes sex-specific deficits in recognition learning and memory and social behavior that persists into adulthood, long after recovery from neonatal anemia (17).

In this study we hypothesized that neonatal PIA acutely impairs mitochondrial respiratory capacity and ETC complex function in the developing hippocampus compared to non-bled control littermates.

## Materials and Methods

### Phlebotomy-induced anemia (PIA) model

The PIA model used in this experiment (**Figure 1A**) has been previously published (15–18). Briefly, beginning at P3, mice were phlebotomized twice daily (5.25 µL/g body weight) until a target threshold of 25% hematocrit was reached. Once at threshold, mice were bled once daily (3.5 µL/g body weight) to maintain the desired degree of anemia throughout the experiment until tissue collection at P14 (n=10 per group). Dams were fed a standard diet containing 200 ppm iron during both pregnancy and lactation.

**Figure 1:**
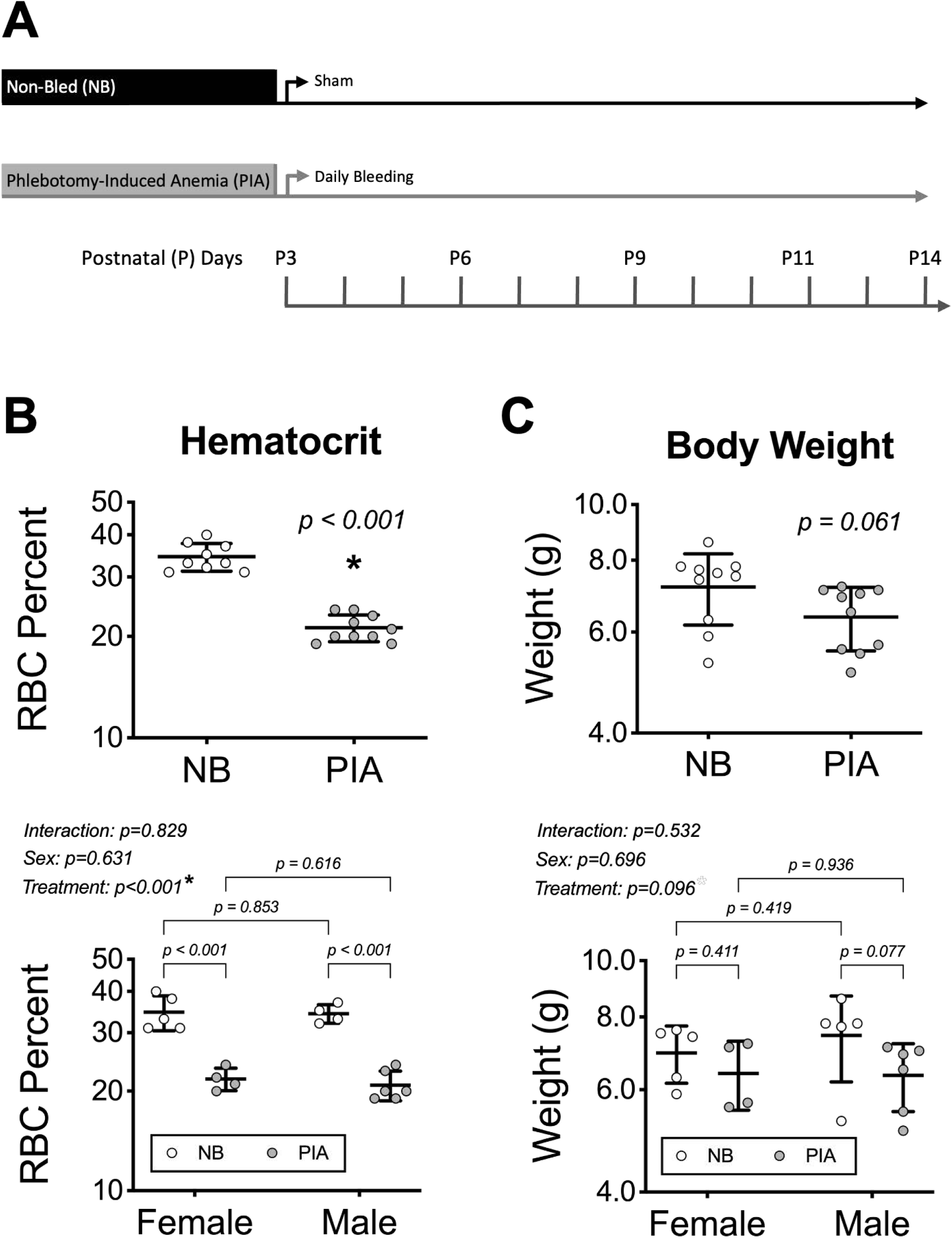
Experimental Design and PIA Pup Characteristics. **A)** To generate neonatal PIA, postnatal day 3 (P3) male and female mice were phlebotomized from the facial vein 2x daily. Once the target threshold of 25% was reached, PIA mice were phlebotomized 1x daily. Non-bled mice received a sham facial needle poke. **B)** Hematocrit (RBC percent of total blood volume) and **C)** body weight was measured at P14 for non-bled (n=9; n=5 female, n=5 male) and PIA (n=10; n=4 female, n=6 male) mice. Bar graphs show all individual values with mean ± SD. Note: P14 hematocrit data was not obtained for one male non-bled mouse. Aggregate male and female data were analyzed by unpaired t-test and p-values are shown for each comparison. Data separated by sex were analyzed by two-way ANOVA and Fisher’s LSD test with p-values shown for individual comparisons and group and interaction effects for treatment and sex. An asterisk (*), if present, indicates a statistically significant effect. NB, non-bled; PIA, phlebotomy-induced anemia; RBC, red blood cell.

### Mitochondrial isolation

After mice were euthanized, the brain was removed from the skull and placed atop filter paper on an ice block. After rapid dissection of the hippocampus, mitochondria were isolated using the Qproteome Mitochondria Isolation Kit (Qiagen; Hilden, Germany) according to the manufacturer’s instructions. Briefly, the tissues were placed in a lysis buffer and homogenized using a pellet pestle (Thermo Fisher Scientific; Waltham, MA) to disrupt the plasma membrane and isolate the cytosolic proteins. The membrane and organelles remained intact and were centrifuged at 1000 x g for 10 min. The resulting pellet was then resuspended in a disruption buffer and passed through a 23-gauge blunt needle to complete cell disruption. When centrifuged (1000 x g for 10 min), the supernatant contains mitochondria and the microsomal fraction. This was recentrifuged at 6000 x g for 10 minutes to produce the mitochondrial pellet. The pellet was then washed and resuspended in a mitochondria storage buffer. Mitochondrial yield from P14 mouse tissue was 100 µg protein per hippocampus on average (range 54-141 µg) as determined by Bradford assay following the isolations.

### Seahorse assays

“Coupling” and “Electron Flow” oxygen consumption rate (OCR) assays were prepared as described (31). All assays were performed with the XFe24 Extracellular Flux Analyzer (Agilent Technologies, Santa Clara, CA). Isolated mitochondria were diluted to 0.1 µg/µl in Mitochondrial Assay Solution (MAS, 1X): 70 mM sucrose, 220 mM mannitol, 10 mM KH2PO4, 5 mM MgCl2, 2 mM HEPES, 1 mM EGTA and 0.2% (w/v) fatty acid-free BSA, pH 7.2 at 37C containing initial substrates (see below) using a 2x MAS solution. Then 5 µg (in 50 µl) was plated. Three to four technical replicate wells were used for each mitochondrial prep from an individual mouse. Three to four background correction wells contained 1x MAS with no mitochondria. The plate was then centrifuged at 2000 g for 20 minutes at 4°C to attach mitochondria to the well bottom. The supernatant was aspirated and 450 µl 1x MAS was added to each well. The mitochondria were examined under a microscope for consistent adherence, then placed at 37°C in room air for 10 minutes.

### Mitochondrial coupling assay experimental design (Table 1)

In the “coupling” assay (31), the initial TCA cycle substrate was 10 mM succinate (ETC complex II specific substrate) with 2 µM rotenone (ETC complex I inhibitor). Injection ports had the following solutions to be released sequentially: 50 µl of 40 mM ADP (4 mM final), 55 µl of 25 µg/ml oligomycin (2.5 µg/ml final), 62 µl of 40 µM FCCP (4 µM final) and 69 µL of 40 µM antimycin A (4 µM final). The following mitochondrial states were measured: **state 2** - basal respiration with substrate present, **state 3** - phosphorylating respiration in presence of ADP and substrate, **state 4o** - non-phosphorylating respiration in the presence of oligomycin (ATP synthase inhibitor), and **state 3u** - maximal uncoupled respiration in the presence of FCCP (disrupts proton gradient and uncouples ETC activity from ATP production). The respiratory control ratio (RCR) was calculated as state 3u/state 4o. State 2 was calculated as the mean of the second baseline measurement. State 3 was calculated as the maximum OCR after ADP injection. State 4o was calculated as the minimum OCR after oligomycin injection. State 3u was calculated as the maximum OCR after FCCP injection. Non-mitochondrial residual respiration was calculated as the minimum OCR value after antimycin A (ETC complex III inhibitor) injection.

**Table 1:** Coupling Assay Seahorse Protocol.

| Command | Time (min) | Drug Injection | Cycles |
| --- | --- | --- | --- |
| Calibrate |  |  |  |
| Mix | 1:00 | None | 2 |
| Wait | 3:00 |  |  |
| Mix | 1:00 | None (Baseline) | 2 |
| Measure | 3:00 |  |  |
| Mix | 0:30 | None | 1 |
| Mix | 0:30 | ADP | 1 |
| Measure | 3:00 |  |  |
| Mix | 1:00 | None | 1 |
| Mix | 0:30 | Oligomycin | 1 |
| Measure | 3:00 |  |  |
| Mix | 1:00 | None | 1 |
| Mix | 0:30 | FCCP | 1 |
| Measure | 3:00 |  |  |
| Mix | 1:00 | None | 1 |
| Mix | 0:30 | Antimycin A | 1 |
| Measure | 3:00 |  |  |

Mitochondrial electron flow assay experimental design (Table 2):

**Table 1:**
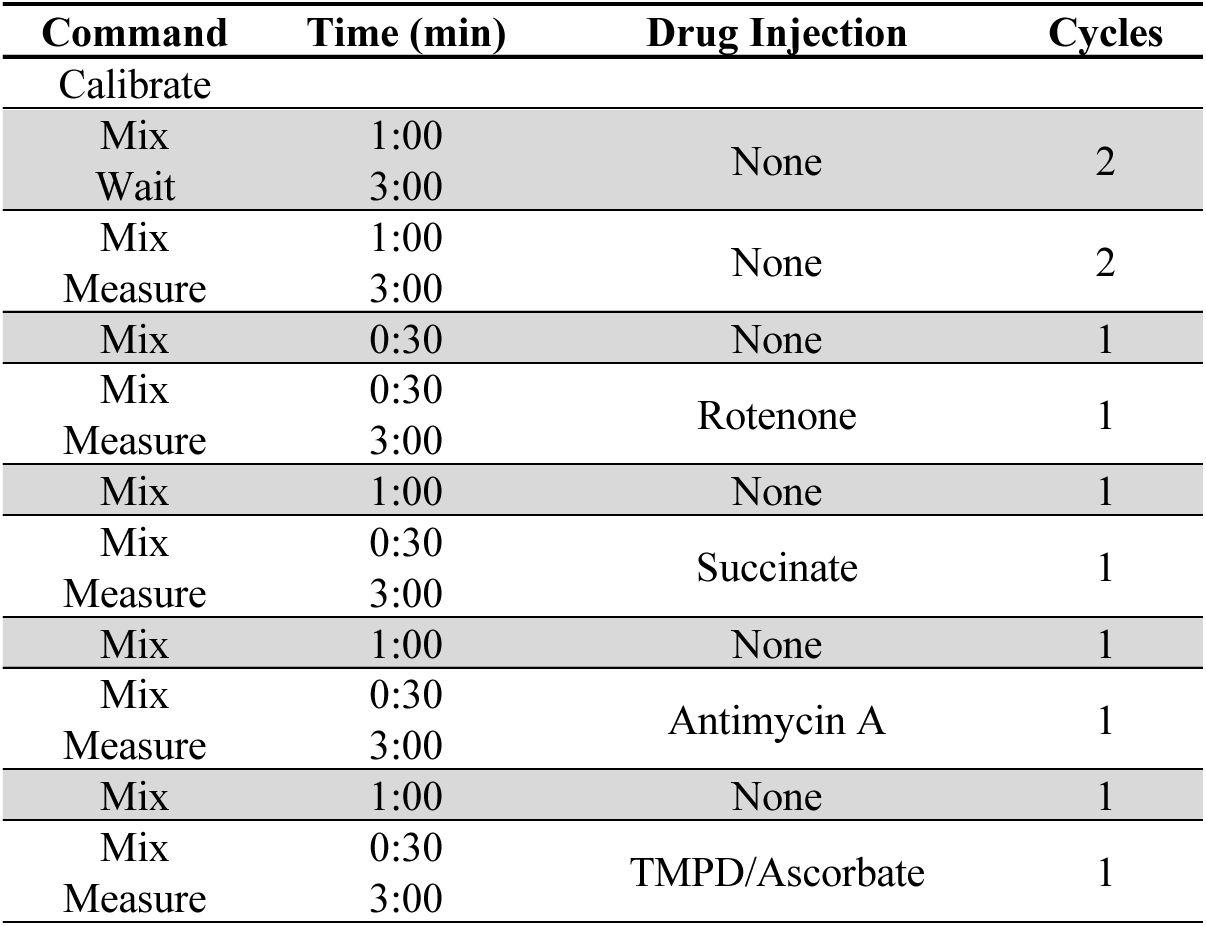
Electron Flow Assay Seahorse Protocol.

In the “electron flow” assay (31), initial TCA cycle substrates were 10 mM pyruvate and 2 mM malate in the presence of 4 µM FCCP () with the following injections in sequential order within the experiment: 50 µl of 20 µM rotenone (2 µM final), 55 µl of 100 mM succinate (10 mM final), 62 µl of 40 µM antimycin A (4 µM final) and 69 µL of 100 mM ascorbate plus 1 mM N,N,N9,N9-tetramethylp-phenylenediamine (TMPD) (10mM and 100 µM final, respectively). ETC complex activities were measured as follows: **complex I** - maximal pyruvate/malate-induced baseline respiration minus the average rotenone-inhibited respiration, **complex II** – maximal succinate-induced respiration minus the average rotenone-inhibited respiration, **complex III** – maximal succinate-induced respiration minus the minimum Antimycin A-inhibited respiration, **complex IV** – maximal TMPD/ascorbate-induced respiration minus the minimum Antimycin A-inhibited respiration.

### Data analysis

Point-to-point data were displayed and analyzed using the Seahorse XF Wave software. Wells that did not respond appropriately to drug injection (e.g., no response or increase after rotenone or Antimycin A) or that had negative OCR values were removed from further analysis. Data from the remaining technical replicates were averaged for each time point. Within each litter, three to four littermates were used per group. Relative OCR values for each mitochondrial state were calculated as a ratio relative to the average non-bled OCR for that state (for each litter). Data from three litters (n=9-10 mice per treatment group) were combined. Statistically significant differences were determined for each mitochondrial state using Student’s t-test (α=0.05). To determine whether sex differences contributed, two-factor (sex and treatment group) ANOVA with Fisher’s least significant difference (LSD) test. Statistical analyses were performed with Graphpad Prism version 10.6 software.

## Results

*Hematology*: Phlebotomy from P3 through P14 reduced packed RBC density to a 21% hematocrit in P14 PIA mice (Figure 1B), a 40% reduction compared to NB controls, which corresponds to a degree of anemia commonly found in preterm infants (4,5,10,11). PIA had a similar effect on the hematocrits of male and female mice. P14 PIA mice also showed a trend towards reduced body weight that was not statistically significant and did not significantly differ by sex (Figure 1C).

*Mitochondrial Coupling Assay*: The coupling assay is used to determine the degree of coupling between the ETC and OXPHOS, and thus efficiency of ATP generation (Figure 2A). Complex II-driven basal mitochondrial respiration, measured and referred to as State 2, was 13% lower (p=0.021) in the P14 PIA hippocampus compared to the hippocampus of non-bled mice (Figure 2B). Adding excess ADP stimulates ATP synthase to produce ATP at maximum capacity (State 3). State 3 was 12% lower (p=0.039) in PIA mice (Figure 2B). Adding oligomycin to inhibit ATP synthase results in State 4o, or non-ADP-stimulated respiration, which is limited by proton re-entry or leak (32). State 4o was 15% lower (p=0.026) in the PIA hippocampus (Figure 2B). Uncoupling of electron transport from OXPHOS by adding FCCP results in state 3u, whereby ETC oxygen consumption is not limited by either ATP synthase or proton re-entry. State 3u was 14% lower (p=0.017) in the PIA hippocampus (Figure 2B). Residual non-ETC mediated respiration, as determined after blocking the ETC with Antimycin A and Rotenone, trended lower (p=0.064) in the PIA hippocampus (Figure 2B). When separated by sex, two-way ANOVA showed a group effect of PIA treatment for States 2, 4o, and 3u, and residual respiration with a trend towards a PIA treatment effect for State 3 (Figure 2C). The only mitochondrial measure with a clear sex-specific effect was residual respiration, which showed a significant difference between non-bled and PIA groups for female mice (p=0.024), but not for male mice (p=0.649). State 3u was significantly reduced in male mice (p=0.047) and not female mice but did show a trend towards reduction in females (p=0.095).

**Figure 2:**
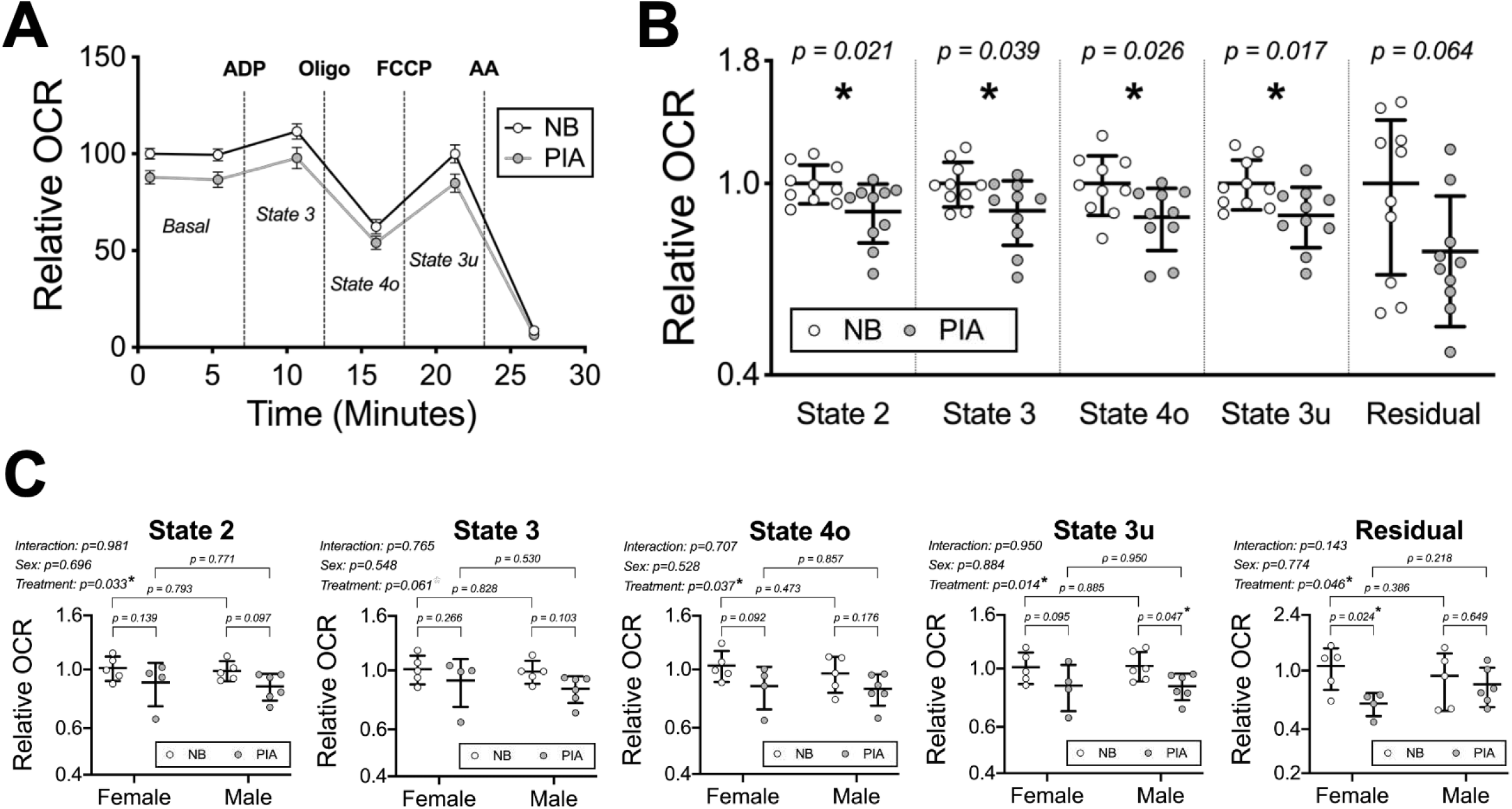
Neonatal PIA reduces hippocampal mitochondrial oxygen consumption rate across functional states. Mitochondria were isolated from the hippocampus of postnatal day 14 (P14) non-bled (n=9; n=5 female, n=5 male) and PIA (n=10; n=4 female, n=6 male) mice. **A)** the mitochondrial coupling assay was performed using Seahorse extracellular flux analysis and mitochondrial OCR for key mitochondrial states was determined. Relative OCR was calculated for each data point relative to the average of the first baseline measurement of the non-bled controls within each litter. X-Y relative OCR data are presented as mean ± SEM rather than ± SD for easier visualization. **B)** Aggregate male and female data for each mitochondrial state were analyzed by unpaired t-test and p-values are shown for each comparison. **C)** Data separated by sex were analyzed for each mitochondrial state by two-way ANOVA and Fisher’s LSD test with p-values shown for individual comparisons and group and interaction effects for treatment and sex. Bar graphs show all individual values with mean ± SD. An asterisk (*), if present, indicates a statistically significant effect. NB, non-bled; PIA, phlebotomy-induced anemia; OCR, oxygen consumption rate.

Since non-ETC mediated residual respiration was lower in the PIA mice, we determined the specific effect of PIA on ETC mitochondrial function by subtracting residual respiration. Basal respiration (p=0.032), maximal respiration (p=0.039), and proton leak (p=0.045) were all significantly lower in hippocampal mitochondria from PIA mice compared to non-bled mice (Figure 3A). Mitochondrial respiration linked to ATP production was not significantly altered in the PIA hippocampus (p=0.165). The RCR indicates how closely coupled the ETC oxygen consumption rate is to mitochondrial ATP production capacity (32). No differences were found between groups for RCR (p=0.880). When separated by sex, two-way ANOVA showed a significant group effect of PIA treatment for basal and maximal respiration with a trend towards significance for proton leak (Figure 3B). There were no significant group effects for sex and no individual comparisons that reached significance.

**Figure 3:**
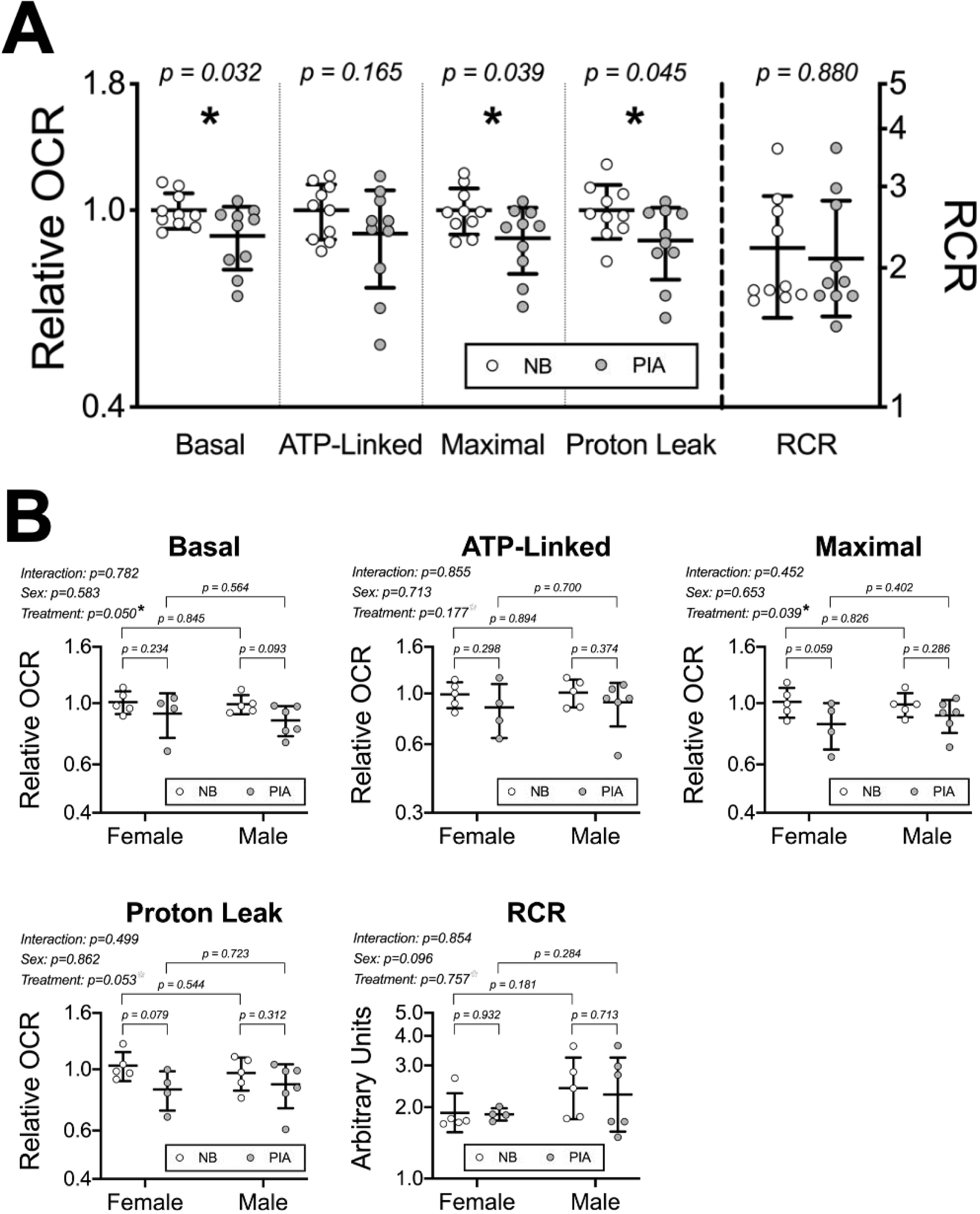
Neonatal PIA reduces key parameters of mitochondrial respiratory capacity in the developing hippocampus. Mitochondria were isolated from the hippocampus of postnatal day 14 (P14) non-bled (n=9; n=5 female, n=5 male) and PIA (n=10; n=4 female, n=6 male) mice. **A)** OCR for key mitochondrial functional parameters (basal, ATP-linked, and maximal respiration and RCR) were calculated from the mitochondrial coupling assay data shown in Figure 2. For each parameter, relative OCR was calculated for each data point relative to the average of the non-bled controls within each litter. Aggregate male and female data for each parameter were analyzed by unpaired t-test and p-values are shown for each comparison. **B)** Data separated by sex were analyzed for each mitochondrial parameter by two-way ANOVA and Fisher’s LSD test with p-values shown for individual comparisons and group and interaction effects for treatment and sex. Bar graphs show all individual values with mean ± SD. An asterisk (*), if present, indicates a statistically significant effect. NB, non-bled; PIA, phlebotomy-induced anemia; OCR, oxygen consumption rate; RCR, respiratory control ratio.

*Mitochondrial electron flow assay*: The electron flow assay sequentially examines the function (i.e. contribution to overall OCR) of the different ETC complexes using pyruvate and malate as TCA cycle substrates to produce NADH for complex I (Figure 4A). Initially, ATP synthase is inhibited with FCCP, so all subsequent steps occur in this uncoupled state. PIA reduced hippocampal mitochondria complex I mediated respiration by 23% (p=0.031) compared to in non-bled mice (Figure 4B). No statistically significant differences were found in complex II, III, or IV mediated respiration between PIA and non-bled hippocampal mitochondria. When separated by sex, two-way ANOVA showed a significant group effect of PIA treatment for complex I-driven respiration with no significant treatment effects for the other complexes (Figure 4C). There were no significant group effects for sex and no individual comparisons that reached significance for any of the ETC complexes.

**Figure 4:**
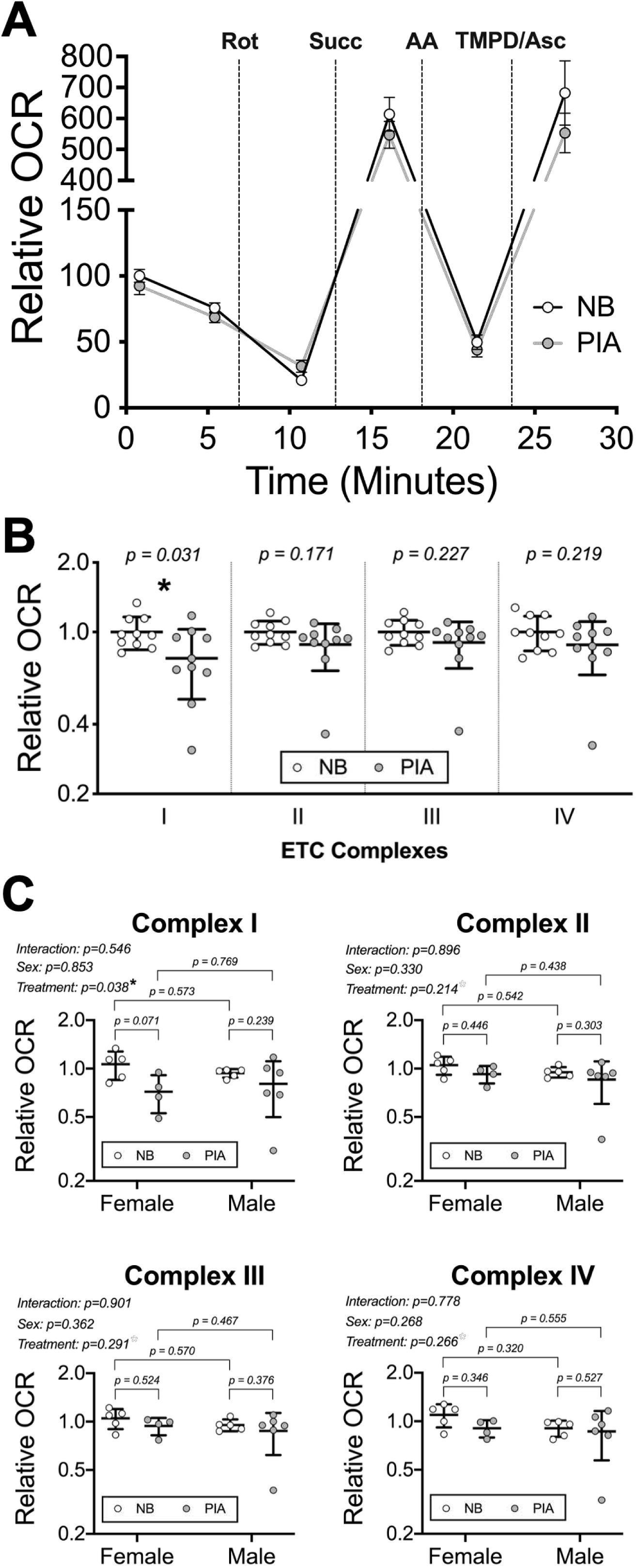
Effect of neonatal PIA on ETC complex functional capacity. Mitochondria were isolated from the hippocampus of postnatal day 14 (P14) non-bled (n=9; n=5 female, n=5 male) and PIA (n=10; n=4 female, n=6 male) mice. **A)** The mitochondrial electron flow assay was performed using Seahorse extracellular flux analysis and mitochondrial OCR for each ETC complex was determined. Relative OCR was calculated for each data point relative to the average of the first baseline measurement of the non-bled controls within each litter. X-Y relative OCR data are presented as mean ± SEM rather than ± SD for easier visualization. **B)** For each ETC complex, relative OCR was calculated for each data point relative to the average of the non-bled controls within each litter. Aggregate male and female data for each parameter were analyzed by unpaired t-test and p-values are shown for each comparison. **C)** Data separated by sex were analyzed for each ETC complex by two-way ANOVA and Fisher’s LSD test with p-values shown for individual comparisons and group and interaction effects for treatment and sex. Bar graphs show all individual values with mean ± SD. An asterisk (*), if present, indicates a statistically significant effect. NB, non-bled; PIA, phlebotomy-induced anemia; OCR, oxygen consumption rate; ETC, electron transport chain.

## Discussion

The major novel finding of this study is that neonatal PIA impairs mitochondrial OXPHOS capacity in the developing mouse hippocampus. This PIA mouse model mimics the degree and timing of anemia of prematurity/phlebotomy (5), a common condition in the neonatal intensive care unit with its etiology closely linked to phlebotomy induced blood loss, during a period of hippocampal development that is similar to the third trimester and early neonatal life in humans (33–35). The adult human brain consumes 20% of the body’s energy despite being only 2% of the body’s weight. The metabolic demands of the developing neonatal human brain are even greater relative to body weight, consuming >60% of the body’s oxygen and glucose (19,20). The neonatal hippocampus, in particular, is undergoing a period of rapid, metabolically demanding growth characterized by neuronal polarization, axon and dendrite growth/branching, synapse formation/refinement, gliogenesis, and axon myelination (35–37). Therefore, impaired hippocampal mitochondrial function may be detrimental to proper neurocircuit formation and ultimately long-term hippocampal functional capacity. Prior studies using this PIA mouse model demonstrate altered transcriptome, neurochemical profiles (e.g., increased lactate) and AMPK/mTOR signaling in the developing hippocampus (15–18). Pre-weanling PIA mice and adult mice that had recovered from neonatal PIA have impaired performance on tasks indexing recognition memory, social interaction, and anxiety (17). The mechanisms driving these long-term neurobehavioral deficits may include impaired neonatal mitochondrial energy production and a permanent blunting of neuronal structural development and functional capacity, as is seen in animal models of early-life ID (21–25,28,30,38–42).

Mitochondria play a critical role in using oxygen to generate energy in the form of ATP to provide the necessary fuel for critical energy-consuming neurodevelopmental processes such as axon/dendrite growth, synapse formation, and myelination. Mitochondrial ATP production requires iron because cytochrome c and all 4 complexes of the ETC depend on iron-sulfur clusters and/or heme groups for their activity (21). TCA cycle enzymes succinate dehydrogenase and aconitase also require iron for their activity. PIA results in a shortage of both oxygen and iron, leaving the mitochondria to either adapt to this limited availability of critical energetic substrates or undergo degeneration.

Mitochondria from the neonatal PIA mouse hippocampus have a lower OCR at states measured where ATP production remains coupled to oxygen consumption including basal (state 2) and ADP-stimulated (state 3). State 2 respiration is dependent on substrate oxidation capacity, ETC function, and proton leak under a physiological condition of low ATP demand (31,32). State 3 respiration demonstrates the ability of mitochondria to respond to a high demand for new ATP synthesis. PIA hippocampal mitochondria also have a lower OCR when ATP synthase is blocked, and electron flow and oxygen consumption is limited by the leak of protons back across the inner mitochondrial membrane from the intermembrane space to the mitochondrial matrix. Less proton leak, and thus less obligate oxygen consumption to maintain the proton gradient, may simply be indicative of the lower overall energetic activity of PIA hippocampal mitochondria. State 3u, mimicking a state of high energetic demand by allowing the ETC to operate at its maximal capacity (not limited by ATP production and ADP availability), was also impaired by PIA.

Together these findings suggest an overall blunting of mitochondrial energetics in the developing PIA hippocampus. We speculate that these mitochondrial deficits are due to brain ID reducing cytochrome and iron-sulfur cluster concentrations (43) and ETC complex activity (23) in the PIA hippocampus. This hypothesis is supported by previous studies in animal models of early life ID (21). In particular, ID reduces the activity of cytochrome c oxidase (complex IV) in the neonatal rat hippocampus (22,23), which is associated with similar metabolic and functional deficits as are seen in the neonatal PIA model (30,38–42). Although isolated mitochondria have access to normal oxygen concentrations during the bioenergetic assays, tissue-level hypoxia also likely plays an important role in programming the *in situ* functional capacity of the mitochondria (16).

Although this study was focused on mitochondrial ETC and OXPHOS activity, the lower non-ETC, residual OCR observed in the PIA hippocampal mitochondria (especially for females), suggests that other mitochondrial oxygen consuming enzymes are also affected in the PIA hippocampus. In addition to OXPHOS (i.e., cytochrome c oxidase), mitochondria consume oxygen during reactive oxygen species generation, heme biosynthesis, sulfur/cysteine metabolism, fatty acid/lipid metabolism, steroid/cholesterol metabolism, ubiquinone synthesis, monoamine metabolism and other metabolic processes (44–50). Iron-containing mitochondrial monooxygenases (e.g., cytochrome P450s), dioxygenases (e.g., cysteamine dioxygenase), oxidases (e.g., monoamine oxidase) and hydroxylases (e.g., coenzyme Q7 hydroxylase) may be particularly affected during PIA due to brain tissue ID (50–53). Moreover, during tissue hypoxia many oxygen-consuming processes are down-regulated through both hypoxia inducible factor (HIF) and non-HIF mediated regulation (50,54–57). Thus, it is not surprising that we see evidence of impaired mitochondrial oxygen-dependent metabolism beyond OXPHOS.

Based on all 4 complexes of the ETC requiring heme and/or iron-sulfur clusters, we predicted that the reduced mitochondrial energetic capacity observed in the coupling assay would involve a reduction in the activity of all ETC complexes. However, we only observed a significant reduction in complex I activity in the electron flow assay. This assay is performed in the presence of FCCP, which uncouples the ETC from ATP synthesis by disrupting the proton gradient across the inner mitochondrial membrane, creating an artificial demand for ETC activity to restore the gradient. Our findings that maximal OCR driven through complexes II, III, and IV is not altered, despite tissue level ID and hypoxia (15,16), suggest that the ETC contains sufficient iron to maintain normal functional capacity in the neonatal PIA hippocampus. This indicates that the reduction in mitochondrial OCR seen in the coupling assay may be due to a reprogrammed, lower metabolic set point. The TCA cycle substrates provided to the mitochondria in the electron flow assay are pyruvate and malate, which predominately produces NADH for Complex I electron transfer. Thus, it is possible that impaired TCA cycle activity contributes to the observed reduction Complex I-driven OCR. One possible mechanism for this is through tissue hypoxia, which has been shown to reprogram the TCA cycle to a lower activity set point in both CNS and non-CNS tissues/cells (58–60). Consistent with this metabolic reprogramming hypothesis, we previously showed that there are higher concentrations of lactate within the neonatal PIA hippocampus (15), indicative of a shift towards anaerobic metabolism. Future work will need to determine the effect of neonatal PIA on additional metabolic pathways contributing to oxidative and non-oxidative ATP generation in the developing brain including glycolysis, TCA cycle, fatty acid oxidation, ketone oxidation and glutaminolysis.

RCR is calculated from the OCRs during state 3 and state 4o and indicates the tightness of the coupling between respiration and phosphorylation, thus indexing mitochondrial efficiency. There were no differences found in this ratio for the PIA mice as compared to their non-bled littermates, indicating the PIA mitochondria maintain similar coupling of OXPHOS to the ETC. This provides additional support that the PIA hippocampal mitochondria are properly functioning, albeit at a lower-than-normal rate in a limited substrate environment (i.e. low oxygen and iron). Being fully functional but producing less energy than control mice may mean that the PIA mice are unable to keep up with the high energy demands of the rapidly growing hippocampus. In previous studies using primary hippocampal neuron cultures, we found a similar pattern following treatment with an iron chelator to induce ID (26,28). Overall mitochondrial respiration was reduced in nearly all measures, but without affecting the ATP coupling efficiency (26,28). This was coupled with a decrease in dendrite complexity, number of branches, and genes indexing dendritic and synaptic development (26,28). This indicates that the impaired mitochondrial respiratory capacity observed in neonatal PIA mice may contribute to impaired early life and adult hippocampal-facilitated behaviors (17) through disrupted neuronal structural development.

Future work will determine whether the reduced energy production observed during the anemic period persists into adulthood, suggesting a reprogramming of mitochondrial regulation in early life consistent with Developmental Origins of Health and Disease principles, and the underlying mechanisms. Furthermore, it needs to be determined whether any interventions such as relief of anemia by red blood cell transfusion or erythropoiesis stimulating agents can increase energy production in the anemic period and prevent long-lasting metabolic, structural and neurobehavioral compromise.

## Acknowledgements

Grants supporting this research included NIH P01 HL046925 (MKG), R01 HD029421 (MKG), R01 HD094809 (MKG), F32 HD085576 (TWB), T32 HL007062 (TWB), F31 NS089093 (DJW) and F30 HD093285 (AKB).

## Author Contributions

All authors contributed to the design of the study; TWB, DJW, and AKB conducted the experiments and analyzed the data; TWB and DJW wrote the manuscript; All authors read, edited, approved and have responsibility for the final content of the manuscript.

## Abbreviations

PIA – phlebotomy-induced anemia, ID – iron deficiency, IDA – iron deficiency anemia, OCR – oxygen consumption rate.

